# A scientific case for revisiting the embryonic chicken model in biomedical research

**DOI:** 10.1101/2024.11.29.623941

**Authors:** Mike J McGrew, Tana Holmes, Megan G Davey

## Abstract

The availability of fertilised chicken eggs and the accessibility and rapid development of the avian embryo, have been utilised in biomedical scientific research to make fundamental discoveries including of developmental processes that are common to all vertebrates, advances in teratology, the understanding of tumour growth and metastasis, angiogenesis, cancer drug assessment and vaccine development as well as advances in understanding avian specific biology. However, recent innovations in chicken transgenesis, genome engineering and surrogate host technology in chickens have only been utilised in a few of these fields of research, specifically some areas of developmental biology, avian sex determination and immunology. To understand why other biomedical fields have not adopted modern chicken transgenic tools, we investigated the non-technical summaries of projects granted in the UK under the Animals (Scientific Procedures) Act 1986 between 2017-2023 to assess when and how chicken embryos are used in research, and if they were considered as a Replacement model for other species.

**Highlights:**

- Evidence for a lack of uptake of the chicken embryo as a partial replacement model
- Case examples of the reduction in animal numbers when chicken models are used
- Proposed action plan for the avian developmental biology community

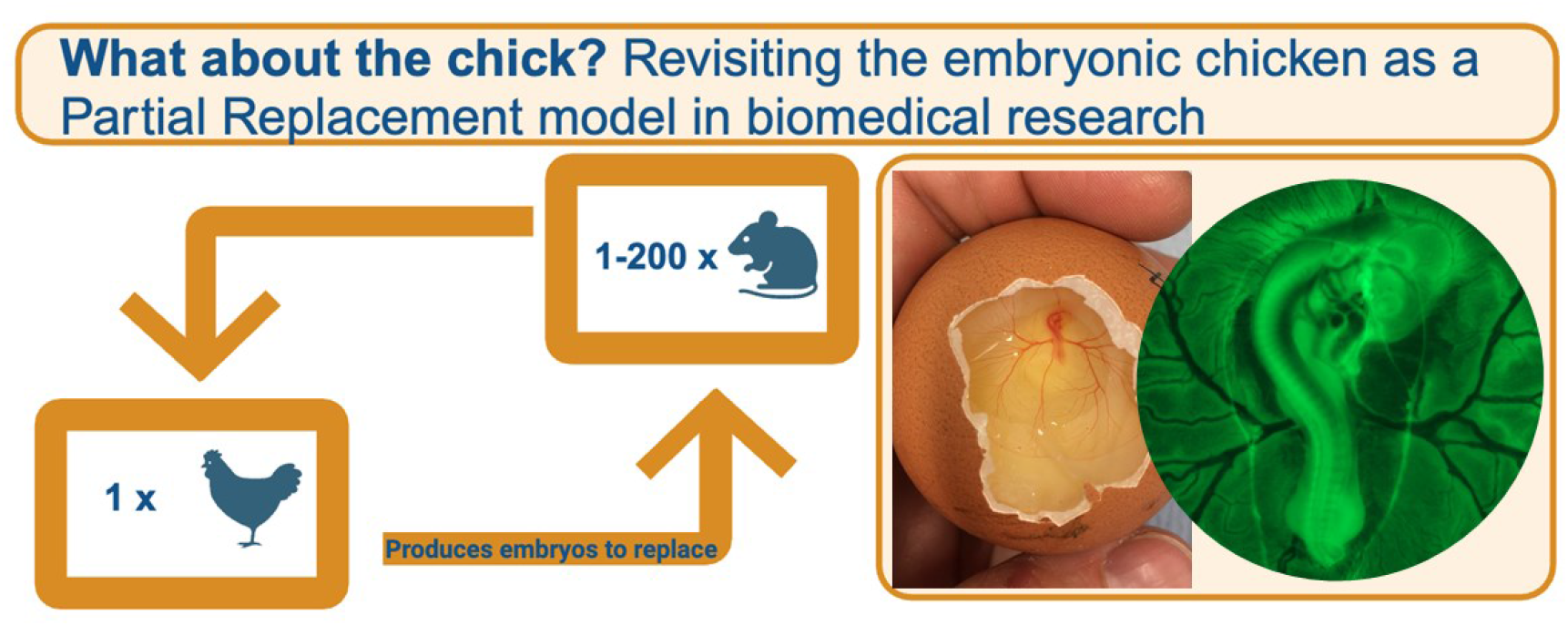

## Introduction

The chicken embryo is an important model for research into the aetiology and treatment of human disease^1^. In the UK, chicken embryos are considered ‘immature vertebrates’ until day 14 of development/incubation are therefore non-protected animals under the Animals (Scientific Procedures) Act 1986. Their experimental use, can therefore replace other protected model organisms, such as mice, in a wide range of biomedical fields, reducing the number of animals used for research^1,2,3^. In addition to research into avian biology and vaccine development, use of the chicken embryo has had significant impact within studies of developmental biology, teratology, animal tumour growth and metastasis, cancer drug assessment and angiogenesis^3,4,5,6,7^.

There is, however, a global failure in uptake of the chicken embryo as a partial replacement model in many areas of biomedical research. Moreover, transgenic chicken resources, such as the Roslin Green eGFP^12^ chicken available to all UK researchers through the National Avian Research Facility (NARF) at The Roslin Institute, and from Clemson University, USA have largely only been adopted by a limited group of developmental biologists. Recent innovations in genome engineering^7,8,9,10^ remain largely underutilised, although they represent a powerful potential for animal Reduction and partial Replacement of several animal models. The utilisation of the chicken embryo chorioallantoic membrane (CAM) assay, is of particular note. The CAM is a highly vascularised and accessible medium for culturing tissues from heterologous species (Fig.1A,B), providing an intermediate model between *in vitro* analysis and *in vivo* animal models^6,7^, in the study of angiogenesis, cancer biology and virology but also in stem cell research, tissue engineering and toxicology^6,7,13,14^. This could be a particular powerful Replacement model as it is inexpensive, requires no specialist equipment, has few ethical considerations and can provide live imaging analysis of processes of interest. Our experience, at The Roslin Institute/National Avian Research Centre (NARF), the only provider of transgenic chicken eggs in the UK, is that transgenic chicken resources are rarely used in the CAM assay, although use of current lines, or development of CAM specific transgenic and bird lines could benefit current CAM users. Other than the CAM assay, in recent years genome engineering technology in chickens, has allowed the generation of knock-out alleles in chickens^9^, providing an alternative route to studies in mouse models to study human genetic diseases. While this might seem novel recent advance, naturally occurring chicken mutants have been used to study human genetic disorders such as polydactyly for many decades^4,5^, and are particularly useful to study mutations which cause early embryonic lethality in mouse models such as ciliopathies^4,5^. This technology, however, has not been widely adopted.

**Figure 1.**
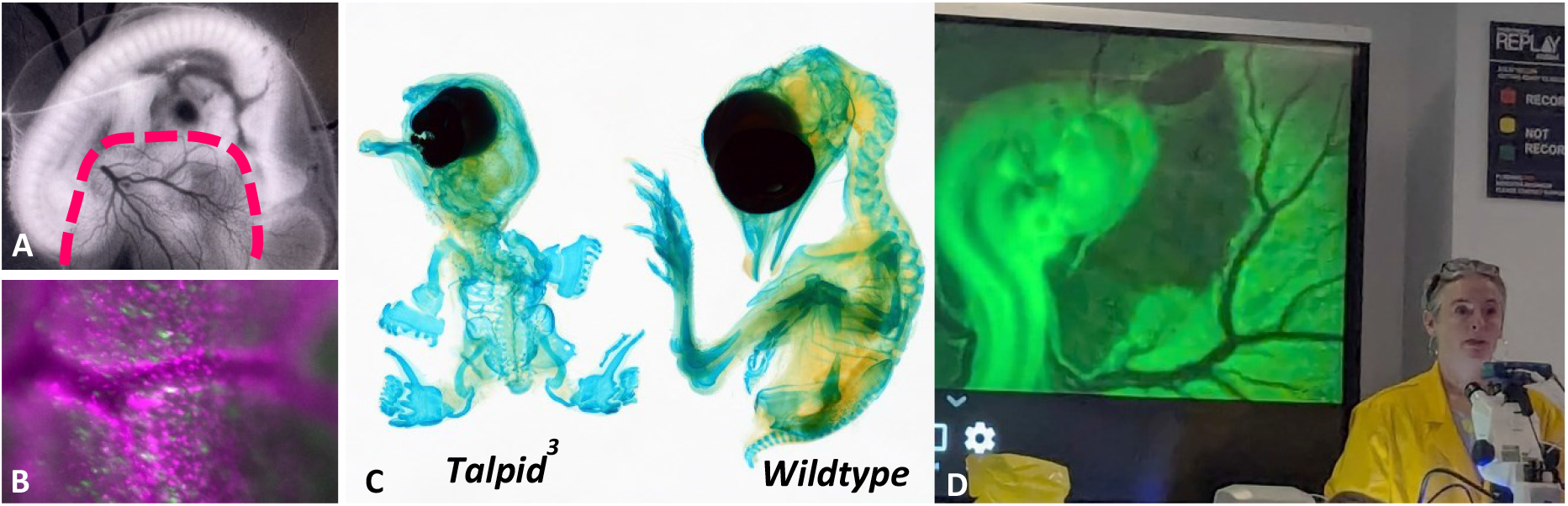
**A**. An eGFP chicken embryo stage 26HH, in greyscale, showing development of the chorioallantoic membrane (pink dashed line). **B**. A newly developed nuclear fluorescent cell cycle reporter, showing expression in the CAM **C**. The *talpid*^*3*^ chicken ciliopathy mutant is a model for the ciliopathy Joubert Syndrome 23. **D**. The Edinburgh Gallus Genomic and Embryonic Development (EGGED) Workshop, held in 2022 and 2024 is a practical workshop to help bridge the translational gaps in chicken transgenesis, developmental biology approaches and the CAM assay. The author, MGD, in yolk yellow coat demonstrating the use of an eGFP transgenic embryo.

In the United Kingdom (UK), the use of regulated under the Animals (Scientific Procedures) Act 1986 (ASPA). ASPA is implemented by the Home Office in England, Scotland and Wales and by the Department for Health, Social Security and Public Safety in Northern Ireland. Researchers wanting to undertake basic or applied research which uses animals, or uses animals in other ‘testing’ or research scenarios, for example testing the quality, effectiveness and safety or drugs or foodstuffs, must apply for a Project Licence. A Project Licence application contains Non-technical summaries which contain a wealth of information on each Project Licence, including the species, number of animals, purpose and type of research, keywords and importantly detailed 3R’s information, including details of how the proposed project has been designed to Replace animals where possible. A recent updated definition of ‘Replacement’ includes distinct definitions of what constitutes a ‘full’ replacement method (Non Animal Method; NAM) or a ‘partial’ replacement model. We suggest there is a potential for improved uptake of the chicken embryo and CAM model in biomedical research as a partial replacement model and that the chicken embryo and CAM assay has not reached its potential as a powerful partial replacement model in biomedical research. To examine this hypothesis, we undertook an analysis of the ‘Application of the 3Rs’ section in Non-technical summaries of Animal Project licenses issued between 2017-2023, as use of chicken embryos or the CAM as a Replacement model should be mentioned here, in projects which have considered their use. Our finding indicate that there is a considerable lack of understanding and uptake of the role the chicken embryo or the CAM assay as a partial replacement models in animal research.

## Materials and Methods

### ‘Non-technical summary’ analysis of Project Licences grants in the UK 2017-2024

‘Non-technical summaries’ for the proposed use of Animals are published as a legal requirement under Article 43 of EU Directive 2010/63 and are available as downloadable PDFs at https://www.gov.uk/government/collections/animals-in-science-regulation-unit. Within Adobe Acrobat, we searched the title and keyword section of 3532 Non-technical summaries of Project licenses issued between 2017-2023. If Non-technical summaries did not contain a Keyword section we also searched the Aims section. In addition we searched for the terms ‘chick’, ‘chicken’, ‘avian’, ‘CAM’, ‘chorioallantois’ with the main body text of all 3532 Non-technical summaries. To discover Project licenses issued to undertake angiogenesis assays we used the search terms: angiogenesis, cancer, cardiovascular, tumour; for Project licenses issued to study ciliopathies with search terms: cilia, cilium, ciliopathy, Joubert, TALPID*; for Project licenses issued to use chicken and CAM models using the search terms: avian, CAM, chicken, chick, chorioallantois. On reading the Non-technical summaries containing search terms of interest, we discounted any studies that were for the purpose of specifically studying avian biology or which used avian embryos for established avian specific protocols, such as, inoculation with influenza virus. We assessed the remaining studies for the inclusion or non-inclusion of the chicken embryo or CAM assay as a partial replacement model.

## Results

Our analysis of Non-technical summaries found that the potential use of the chicken embryo as a partial replacement model were rarely considered.

### Chorioallantoic Membrane Assays

Between 2017-2023, the chicken chorioallantoic membrane (CAM) assay was referenced in only 9 Non-technical summaries, section for Project licences granted under the Animals (Scientific Procedures) Act 1986, for studies of cancer, cancer therapies, blood vessel development or, in one case, biomaterial engineering (Table 1). In comparison 662 Non-technical summaries investigating cancers, tumours, angiogenesis or cardiovascular biology did not consider the use of the CAM or chicken embryo as a partial replacement model. To undertake CAM assays before 14 days of incubation, would not normally require Project Licence and so this does not represent the number of CAM assays undertaken in the UK. It does highlight however, how infrequently the CAM assay is considered as a partial replacement model when designing experiments to investigate cancer, cancer therapies, angiogenesis.

**Table 1.**
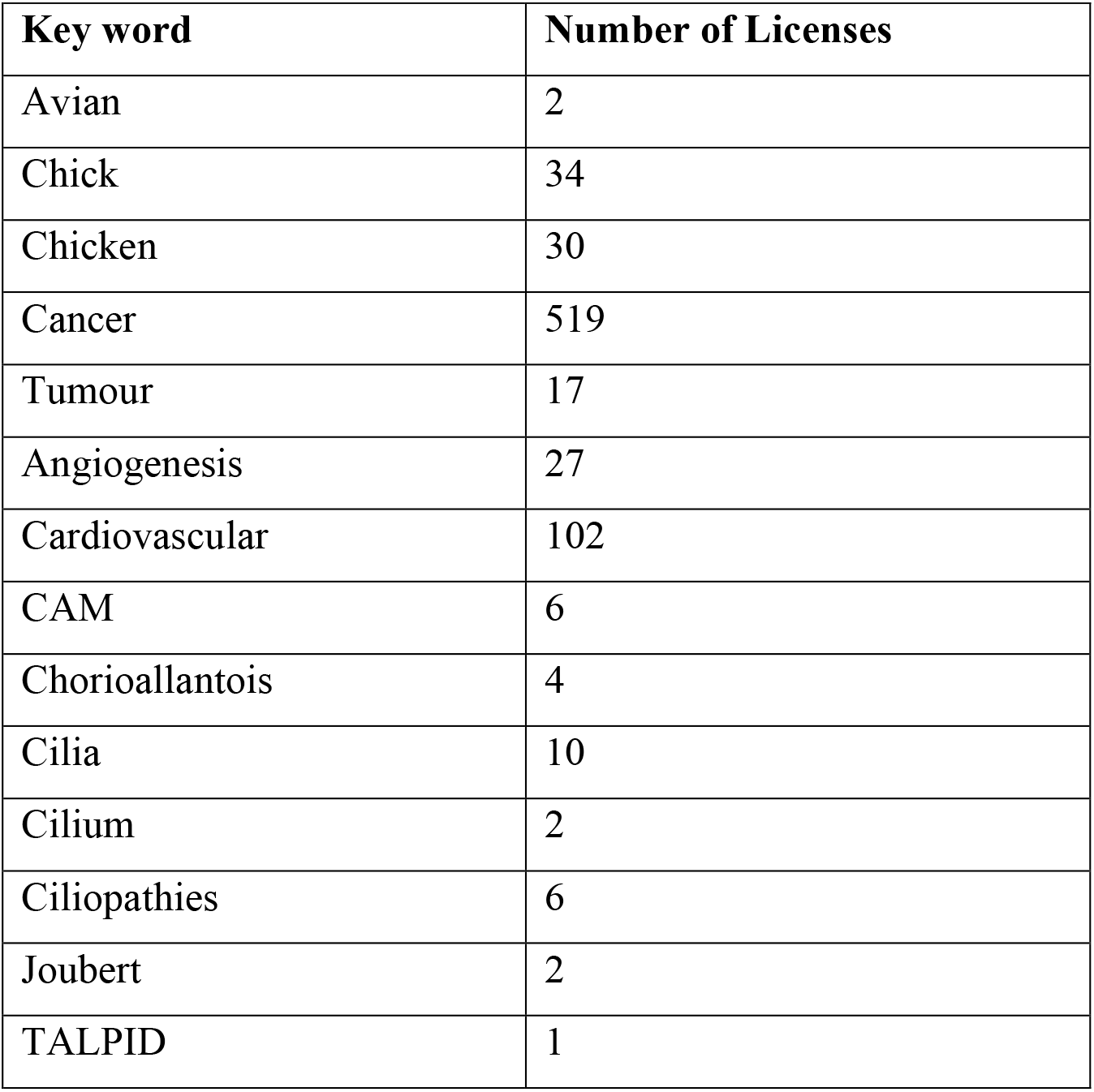
Summary of Keyword Mentions in 3532 Non-technical summaries for Project Licenses Issued 2017-2023.

| Key word | Number of Licenses |
| --- | --- |
| Avian | 2 |
| Chick | 34 |
| Chicken | 30 |
| Cancer | 519 |
| Tumour | 17 |
| Angiogenesis | 27 |
| Cardiovascular | 102 |
| CAM | 6 |
| Chorioallantois | 4 |
| Cilia | 10 |
| Cilium | 2 |
| Ciliopathies | 6 |
| Joubert | 2 |
| TALPID | 1 |

This is highlighted by the very contrasting statements taken from two applications to studying cancer-

> “Within our School we are working on developing the chick chorioallantoic membrane (CAM) as an alternative, high-throughput method for the development of novel cancer imaging agents. The CAM is a highly vascularised extra-embryonic membrane of the chick embryo. The CAM can be accessed easily with minimal invasion to the embryo, enabling the growth of cultured cancer cell and patient-derived xenografts, complete with a co-opted vascular system. This may serve as a replacement for some animal experiments in the future.” (Project 124 2022: In Vivo Imaging and Studies in Cancer Disease Models)
>
> “This complexity cannot be recapitulated in any in vitro system or lower organismal system such as flies or fish. We are not aware of any alternatives to the utilisation of mice for in vivo experiments.” (Project 195 2017: Cancer development, growth, detection & treatment)

Animal Project licenses issued in 2017 and 2018 alone, which included ‘angiogenesis’ in the non-technical summary, included 8 projects^15^ anticipating the use of approximately 135,000 mice over a five year period, as well as many thousand rats^15^, in studies of cancer and tumour development, growth, detection or treatment and cardiovascular disease and remodelling. None of these 8 projects considered the use of the chicken CAM assay in the ‘Application of the 3Rs’ section of the project license application. By comparison, one project (‘Study effects anti-cancer agents on tumours’; Project 16, 2017) proposed to use only the chicken CAM assay in the study of angiogenesis. This project anticipated the use of 2500 chicken embryos over five years, a number easily generated from a small flock of 8 or less birds and demonstrates that it is likely that angiogenesis assays undertaken in mice could be replaced by the CAM assay. If 135,000 angiogenesis assays in mouse could be replaced by 1:1 with chicken embryos, then these could be produced by the use of approximately 675 female birds over 5 years, a 200-fold reduction in animal use. In the UK in 2020, 184,254/827,801 procedures undertaken in mice were for the study of ‘Oncology’, the ‘Cardiovascular Blood and Lymphatic System’, ‘Human Cardiovascular disorders’ or ‘Cancers’ (Annual Statistics of Scientific Procedures on Living Animals, Great Britain 2020), many of which are likely to be angiogenesis assays. We propose that wider uptake of the CAM assay in these fields of research, could significantly reduce the yearly number of procedures undertaken in mice and other rodents.

### The chicken embryo as a 3R biomedical model

The bias against the chicken embryo was not only limited to the CAM and angiogenesis assays. For many hundreds of Project licenses issued between 2017-2023, the case for mouse studies were made citing superior genetics, the ability to undertake genome engineering and imaging approaches and inability to model human anatomy, physiology and diseases in non-mammalian models (Table1).

> “Whilst less sentient animals such as zebrafish can provide some insights into basic aspects of vertebrate development, and are used by research teams to complement mouse studies, to understand conditions affecting such tissues and organs as the kidney, immune system, lungs and brain, a mammal is needed.” (Project 23 2020-Breeding and production of genetically altered mice, 7450 mice).

This specific example, was to undertake a study of Joubert’s syndrome, a recessive ciliopathy syndrome for which the Davey Lab has published 12 research papers and 3 reviews^4,5^, using the *TALPID3*^*-/-*^ chicken embryo to model human anatomy and physiology (Fig.1C). Surprisingly, given that the *TALPID3* ciliopathy locus was discovered in chickens, Project 102 2020: Molecular Basis of Development and Disease, proposed using mice to study “The molecular changes associated with mutations in basal body proteins such as Talpid3 seen in the ciliopathy-Joubert syndrome…” without any consideration using the *talpid*^*3*^ chicken line.

In this instance, *talpid*^*3*^ is an excellent example of how using the chicken as a model can dramatically reduce the number of animals used in research. A flock of *TALPID3*^*+/-*^chickens (2 *TALPID3*^*+/-*^ males and 5 *TALPID3*^*+/-*^ hens), produces 1300 fertile eggs/year, ∼25% are *TALPID3*^*-/-*^. Thus, 7 birds produce approximately 300 *TALPID3*^*-/-*^ embryos which can be used to study a Joubert syndrome phenotype. To obtain the same number of *Talpid3*^*-/-*^ mouse embryos would require ∼170 *Talpid3*^*+/-*^ dams mated with *Talpid3*^*+/-*^ males. To produce this number of *Talpid3*^*+/-*^ mice, would likely require breeding of up to 700 mice. This demonstrates that use of a chicken embryo model to study a recessive human condition, could result in a 100 fold decrease in the number of animals.

These animal numbers are supported by evidence from Project license applications. When chicken embryos are considered within experimental design, they had a significant role in replacing animal models. Of the 6/1062 Project licences issued in 2017 and 2018, which proposed the use of chicken embryos as a partial replacement for animal models, the proportion of chicken embryos used as partial replacement anticipated animals was often high, totalling 63,300 replacement experiments^16^.

### Conclusion and Points for Action for Our Community

#### Barriers to uptake of the CAM assay and the chicken embryo as a model

If the chicken embryo is such an excellent replacement model, why is the current uptake so poor? While the chicken embryo is recognised as a historically important model in developmental biology^1,2,5^ and angiogenesis^6,7,13,14^, its use declined due to the strength of genetic, transgenic and latterly, genome engineering approaches in mice. However recent major advances in transgenic and genome editing technologies in the chicken, have updated and augmented the chicken embryo as a tractable research model^8-12^. We suggest that the knowledge exchange of these advances to the biomedical fields has been lacking.

We recognise that obstacles to use of the chicken embryo include a lack of knowledge exchange and information sharing, between a diverse range of researchers and biomedical fields. There is particularly a general lack of knowledge regarding the following; the ease of uptake and use of the chicken embryo in a ‘mammalian’ focused research setting, the availability and types of transgenic chicken resources, the process to generate new genome engineered models through sterile surrogate host technology, UK legislation regarding work in chicken embryos and the potential and acceptance of avian models to model mammalian, particularly human genetics, anatomy, development and physiology.

While poultry husbandry courses are common and available in most countries, training in use of the chicken egg for developmental biology is extremely limited. Since 2022, The Roslin Institute has developed the ‘EGGED Workshop’, which we hope to run biannually to overcome many of these barriers (Fig.1C, D). We aim to bridge the knowledge gap of non-chicken experts by providing practical experience of chicken embryo use, publicising the transgenic and genome engineering resources available through National Avian Reasearch Facility (NARF), supporting knowledge exchange regarding chicken husbandry and production, Home Office legislation related to chicken embryo work and best practise in Public engagement concerning the use of Animals in Research and 3Rs objectives. In addition as developmental biologists who teach undergraduates, we also see the chicken embryo as a powerful teaching tool, which communicates the usefulness of this model to future scientists. As a community we could develop communal teaching resources that highlight the resourceful chicken model. All these resources could be expanded to underpin knowledge and skills transfer available through EGGED or other training workshops via development of online resources and engagement with a new audience, who could utilise the chicken embryo and CAM model.

## Funding

This study was funded by Biotechnology and Biological Sciences Research Council (BBSRC) to The Roslin Institute, BB/X010937/1 MGD

**Conceptualization:** MGD

**Methodology and resources:** MGD.

**Analysis and investigation:** MGD, TH

**Writing – Original Draft:** MGD, MM. All authors contributed to manuscript review and editing

## Data sources from the following

**https://www.gov.uk/government/collections/animals-in-science-regulation-unit**

